# Effects of bicycle geometry and riding position on the potential of residual limb muscles to pedaling with a transfemoral prosthesis: a computer simulation study

**DOI:** 10.1101/615245

**Authors:** Yusuke Okita, Takashi Nakamura

**Affiliations:** Department of Prosthetics and Orthotics, Research Institute, National Rehabilitation Center for Persons with Disabilities, Tokorozawa, Japan

**Keywords:** transfemoral amputation, cycling, muscle potential, musculoskeletal model, induced acceleration

## Abstract

We performed musculoskeletal simulations to provide information on the effects of riding position and bicycle geometry on pedaling with a transfemoral prosthesis. Sixty-four models and their corresponding kinematics in one pedaling cycle were generated from the baseline one-leg cycling model by varying one of the six variables (seat height, seat-tube angle, crank length, pelvic tilt, anteroposterior seating position, and thigh length relative to the leg). Induced acceleration analysis was performed to compute the potentials of the residual hip muscles for crank rotation in each model. The simulation results quantified the effects of each variable on the hip and knee kinematics and muscle potential during a pedaling cycle; seat height, crank length, and pelvic tilt were the primary candidates for bicycle fitting considering their accessibility and simple effects on the joint kinematics and muscle potential. The seat-tube angle (similar to pelvic tilt) and the anteroposterior seating position (similar to seat height and seat-tube angle) seemed to have an effect similar to the other variables and thus can be reserved for fine-tuning after gross fitting of the bicycle. Although not considered for adjustment, considering the effects of the thigh length could help as it affects hip kinematics and muscle potentials.

## Introduction

Cycling is one of the most challenging activities for individuals with transfemoral amputation (TFA) wearing a passive prosthesis that lacks active control of the knee and ankle joint. Cycling and pedaling exercises could be beneficial for individuals with TFA by providing aerobic conditioning^1,2^, participation in recreational or competitive activities^3^, and efficient transportation. Although individuals with TFA have an option to pedal using the contralateral leg only^2,4^, pedaling with a prosthetic leg could add to the power on the crank and favorably maximize the use of the residual limb muscles during cycling.

Clinicians rely on empirical knowledge in teaching and training for pedaling due to the paucity of biomechanical knowledge on cycling after lower limb amputation even though pedaling has long been investigated in biomechanical studies^5–9^. Previous studies on pedaling after lower limb amputation have mainly focused on transtibial amputation (TTA)^10–13^ and less on TFA.^12^ Childers et al.^12^ reported that the pedal force effectiveness of two subjects with TFA, which can be derived from the radial and tangential pedal force, was similar for those with TTA. However, a movement strategy for pedaling specific to TFA might exist because the physiological knee joint is absent after TFA in contrast to after TTA.

Individuals with TFA must rely on the residual hip muscles to pedal with their prosthetic limb. The gluteus maximus and hamstrings are regarded as the primary hip extensors during pedaling^5,8,14^, but hip adductors and abductors also contribute to the flexion-extension movement of the hip in specific joint angles^15,16^. Hip abductors are one of the target muscles for strengthening in prosthetic rehabilitation^1,17,18^ because of their ability to support and balance the body during locomotion.^19–21^ Hip adductors are less focused on but are known to be activated during pedaling with an intact limb.^22^ The residual hip adductors tend to retain their volume according to the residual limb length,^23^ which is associated with walking performance^24–26^. The reduced number of active joints in the passive prosthetic leg could necessitate a fine-tuned seat configuration, crank length, and riding position for effective pedaling with a prosthetic leg. When individuals with TFA learn to pedal, the bicycle geometry and riding position should be optimized because they are related to the required joint range of motion and the efficiency of pedaling. However, baseline biomechanical information with which clinicians can use to set these configurations to fit the bike to each individual is scarce. Experiments manipulating riding conditions could yield meaningful results, but the number of conditions available during one experimental session is limited because of the fatigue and variability of the limb length and the available muscles after amputation surgery. Due to this fact, computer simulation could be an option for testing multiple conditions. Computer simulation using musculoskeletal models enables one to evaluate individual muscle function such as the contribution of the muscle forces to crank angular acceleration, which is practically impossible to know from an experiment and thus provides a better understanding of pedaling biomechanics for individuals with TFA.

In this study, we quantified and evaluated the effects of bicycle geometry and riding position on pedaling kinematics and muscle potential on pedaling performance using a musculoskeletal simulation. Because of the exploratory nature of this study, we did not prepare specific primary hypotheses. However, we anticipated that the responses in kinematics and muscle potential would depend on the manipulated parameters from which we could suggest how to optimize cycling/pedaling for individuals with a TFA. The target population of this study was transport/recreational cyclists with transfemoral prostheses or individuals who prefer pedaling as an exercise modality for their residual limb muscles and not competitive cyclists who need to maximize power output by making the most of their bilateral leg.

## Materials and Methods

### Study design

This was a computational simulation study without experimentally collected motion data and thus did not require ethical approval. We developed a musculoskeletal model to simulate pedaling using a transfemoral prosthesis and analyzed the potential of individual residual limb muscles to rotate the crank during a pedaling cycle by varying the bicycle geometry and riding position. OpenSim^27^ (version 4.0) and MATLAB (version 2012b, Mathworks, Natick, MA, USA) were used for the whole analysis.

### Pedaling model

A unilateral TFA pedaling model (pedaling model, Figure 1) was created by modifying an open-source musculoskeletal model^28,29^ representing an average-sized male adult (height: 1.70 m, body mass: 75 kg). The original model was reduced and modified to reflect the body configuration of individuals with TFA and to facilitate pedaling simulation. The pedaling model included segments representing the pelvis, thigh, shank-foot, pedal, and crank. The model also included pin joints representing the hip, knee, pedal, and crank center. The pelvis and crank were fixed to the ground such that the seating position was defined by the distance between the hip joint center, the crank axis on the sagittal plane (seat height), the seat-tube angle, and the anteroposterior (AP) seating position. The distance between the foot and the pedal was fixed by a weld constraint. This configuration of the model resulted in a pseudo one degree of freedom, similar to the previous simple pedaling model representing healthy subjects.^30^ However, this model can also represent the pedaling of individuals with TFA who use a passive prosthetic leg with reduced degrees of freedom. The default mediolateral and axial rotation angles of the hip and knee joints were set to zero. Thererfore, the effects of the axial rotation of the femur were not considered. The long head of the biceps femoris, semimembranosus, and semitendinosus were transferred to the distal end of the femur with a via- point assuming a 4 cm diameter of the femoral bone (Figure 1.) The pedaling model represented a person with TFA and a long residual limb or with knee disarticulation. The thigh and shank length of the pedaling model were derived from the original model and were 0.4096 m and 0.4 m, respectively. The distance between the ankle joint and the pedal axis was additionally defined as 0.115 m; therefore, we defined the default total leg length as 0.9246 m.

**Figure 1.**
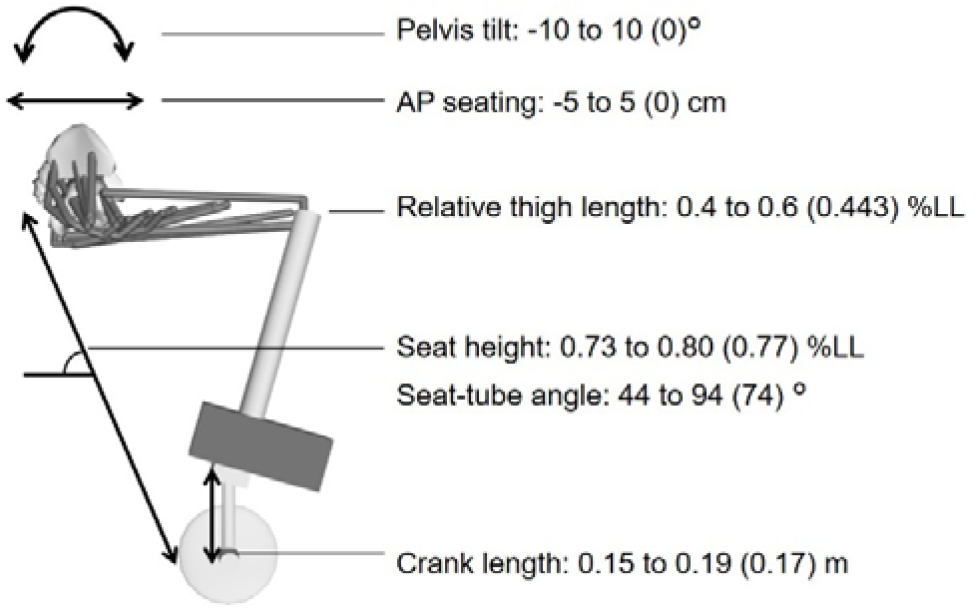
The pedaling model and the parameters manipulated for this study. The range (default value) of each parameter is indicated. AP: anteroposterior, LL: leg length (sum of the femur and shank length and the foot-pedal height)

### Variables and baseline condition

We evaluated the effects of altering each of the six variables representing the bicycle geometry (seat height, seat-tube angle, crank length) and the riding position (pelvic tilt, AP seating position, thigh length relative to total leg length [relative thigh length]). The baseline condition was defined as seat height = 77% leg length, seat-tube angle = 74°, crank length = 170 mm, pelvic tilt = 0°, AP seating position = 0 cm, and relative thigh length = 0.443 (Figure 1.) The seat height for the baseline condition was determined as the median value of preliminary analysis results where pedaling models with 73 to 81 % leg length could be generated. Other baseline values were determined by considering the neutral position of a rider for the riding position, and the geometry of conventional bicycles for the seat-tube angle and crank length. From the baseline model, we generated a total of 64 models by changing each of the six variables at a time to evaluate the effect of changing a single parameter. The range of values for each variable was determined by considering the practically possible range of adjustment (for crank length, pelvic tilt, AP seating position, and relative thigh length) or by preliminary analysis (for seat height and seat-tube angle). On this terms, we explored the range of values for each variable with which the pedaling model could achieve a pedaling cycle by kinematic simulation as described next.

### Generating pedaling kinematics for each model

Forward kinematic simulations were performed to obtain the pedaling kinematics for each model by assembling the model segments at crank angles between 0° and 360° with a 1° increment. The assembly algorithm^31^ searched for the body kinematics that least violated the kinematic constraint at each crank angle using the interior point method. For the model assembly, the pedal and the prosthetic foot were constrained, but slight movement was allowed between them for computational efficiency, while the pelvis and crank were fixed to the ground. A pedaling cycle (0° to 360°) for each condition was regarded as achievable if the maximum distance between the foot and pedal did not exceed 0.3 mm; the threshold was chosen based on our preliminary analysis. Derived kinematic data were used as inputs for the following induced acceleration analysis.

### Computing potential of residual muscles to crank angular acceleration

We evaluated the potentials of the residual hip muscles for angular acceleration of the crank at each crank angle using induced acceleration analysis that computed the angular acceleration of the crank induced by a unit force (1 N) of each residual muscle. A positive value (in rad/s^2^) indicated the potential of the forward angular acceleration of the crank. The results on the hip muscles of interests (Table 1) were presented based on their expected contribution to hip extension during the first half (i.e., downstroke phase) of the pedaling cycle.

**Table 1.**
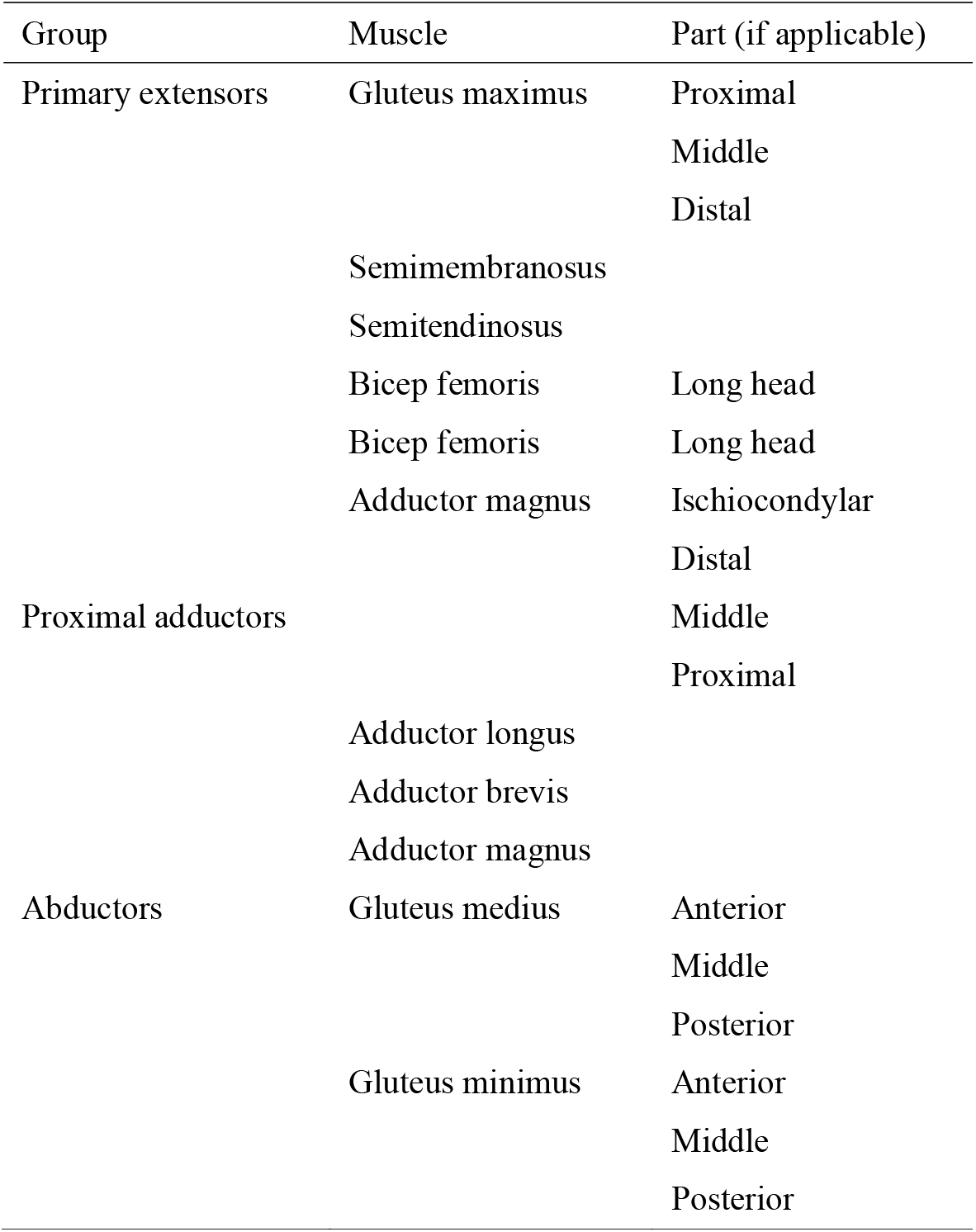
Hip muscles and muscle groups

The effects of each variable on the hip and knee kinematics were evaluated using the values and timings of peak flexion/extension angles and the range of motion during a pedaling cycle. The effects of each variable on individual muscle potentials to rotate the crank forward were evaluated using the maximum value and its timing, and the range of crank angles that each muscle could rotate the crank forward. This study only evaluated pedaling with a passive prosthesis without fixing the foot on the pedal by a cleat, and thus we focused on the capacity of residual hip muscles to rotate the crank forward in the downstroke phase. The results of an entire pedaling cycle were showed to facilitate future studies focusing more on competitive cycling or cycling with a powered prosthesis, although we did not evaluate the upstroke phase where the hip and knee flexors can also contribute to forward crank rotation.

## Results

With regards to hip and knee kinematics, a greater seat height reduced the hip and knee joint angles (Figure 2A). The effects of the seat height were greater on the minimum joint flexion (−2.1° for the hip and −4.1° for the knee per unit height [% leg length] increase), than on the maximum joint flexion (1.1° for the hip and 1.4° for the knee). The hip angle reached its minimum value earlier with a greater seat height than a lower seat height (2.3° earlier with unit height increase). Altering the seat-tube angle induced a corresponding vertical and horizontal shift of the kinematic trajectories for both joints. For a constant range of motion, a greater seat-tube angle induced a delayed peak flex/extension for both joints and smaller hip flexion angles throughout the pedaling cycle. The maximum and minimum knee flexion angle was not affected by changing the seat-tube angle (Figure 2B). The crank length affected the range of motion for the hip and knee. By extending the crank length by 5 mm (more practically implementable than 1 mm), the range of motion was increased by 1.5° for the hip and 2.5° for the knee (Figure 2C). Increasing the anterior pelvic tilt induced a corresponding degree of hip flexion (Figure 2D) without affecting the knee kinematics. The anterior seating position reduced the hip flexion in the upstroke phase (i.e., the leg is in flexion) and increased the knee flexion angles in the downstroke phase (i.e., the leg is in extension) (Figure 2E). As the relative thigh length increased, the range of motion of the hip (1.0° reduction per unit length) and maximum hip flexion angle (1.4° reduction per unit length) was reduced (Figure 2F).

**Figure 2.**
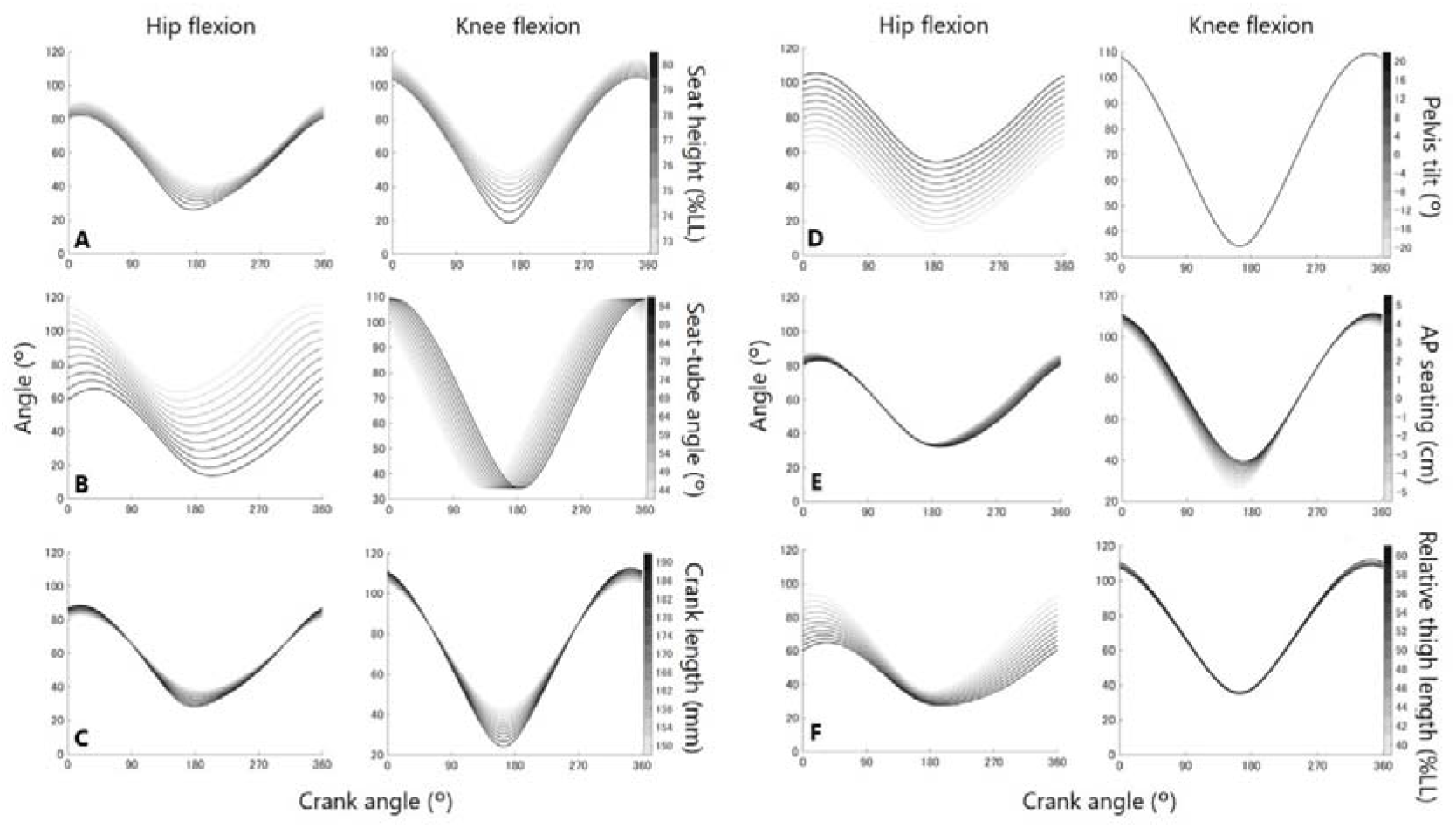
Hip and knee kinematics during a pedaling cycle achieved by each model. The color density of the line indicates the value of the variables representing bicycle geometry or riding position.

With regards to the muscle potential to rotate the crank, individual muscles were categorized to three groups based on the shape of the waveform: primary extensors (gluteus maximus, hamstrings, and the distal and ischiocondylar part of adductor magnus), proximal adductors (adductor brevis, adductor longus, and the proximal and middle part of adductor magnus), and abductors (gluteus medius and gluteus minimus). The seat height, crank length, AP seating position, and relative thigh length affected the range of crank angles that the residual muscles could rotate the crank forward, while the seat-tube angle, pelvic tilt, and relative thigh length affected the peak values or the shape of the waveform (Figure 3–4 and Supplementary figures.) As the seat height increased, the offset of crank acceleration came earlier (Figure 3A). As the seat-tube angle increased, a corresponding amount of horizontal and vertical shift of the waveform was observed for all the muscles evaluated. Increasing the seat-tube angle increased the potential to rotate the crank forward for primary extensors but reduced it for proximal adductor muscles (Figure 3B.) A longer crank length changed the offset with a slight change in peak values (Figure 3C). Tilting the pelvis anteriorly resulted in an increased potential of the proximal adductors and reduced that of other muscles (Figure 4A). Anterior transitioning of the seating position and increasing the length of the thigh relative to the shank caused a delayed onset and offset of forward acceleration of the crank with little change in peak values (Figure 4B and C).

**Figure 3.**
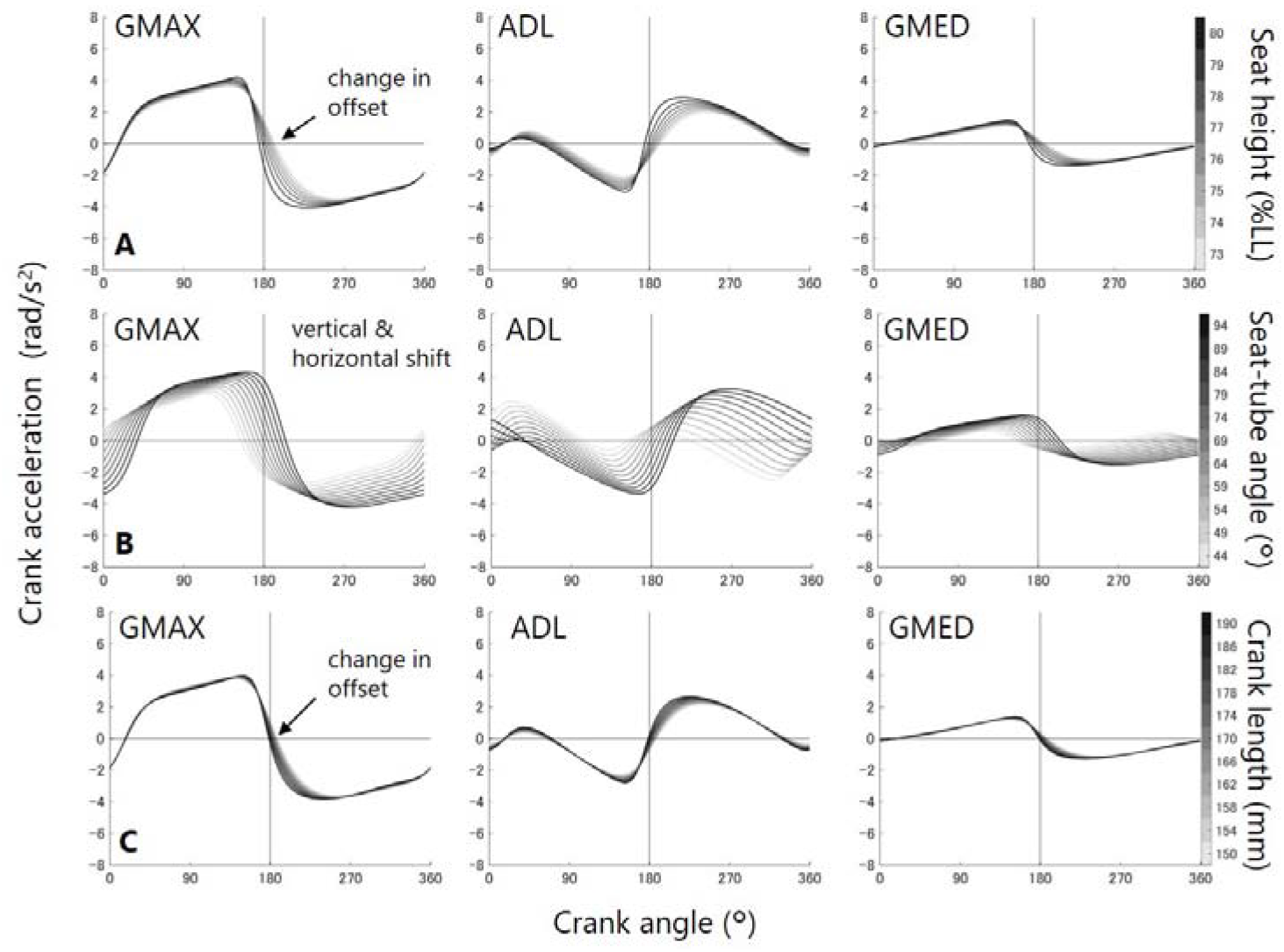
Potential of the residual hip muscles to rotate the crank depending on bicycle geometry (A: seat height, B: seat-tube angle, C: crank length). The middle part of gluteus maximus (GMAX), adductor longus (ADL), and the middle part of gluteus medius (GMED) were presented representing three muscle group (primary extensors, proximal adductors, and abductors, respectively).

## Discussion

In this study, we investigated pedaling kinematics and the residual muscles’ potential to rotate the crank by varying the bicycle geometry and the riding position using a musculoskeletal model and computer simulation. The simulation results quantified the effects of six different variables on the hip and knee kinematics and muscle potential during a pedaling cycle; seat height, crank length, and pelvic tilt were the primary candidates for bicycle fitting considering their accessibility and simple effects on the joint kinematics and muscle potential. The seat-tube angle and the anteroposterior seating position seemed to have an effect similar to the other variables and thus can be reserved for fine-tuning after gross fitting of the bicycle.

Residual hip muscles are the primary actuators for pedaling for people with a transfemoral prosthesis. Thus, information on the potential of these muscles could help optimize bicycle fitting and pedaling performance for those people and lead to a better understanding of the function of the residual hip muscles during movement with a prosthetic leg. The effects of bicycle geometry and riding position on cycling performance have been investigated for healthy subjects ^8,32–34^, but these findings assume the presence of an active knee and ankle movement, which does not apply to individuals with TFA.

The seat height affected not only the range of motion during pedaling but also the potential of residual hip muscles for pedaling, especially at the end of the downstroke phase. The kinematic response to the change in seat height differed between the hip and the knee, possibly because of the difference in the thigh and the shank-foot length. Increasing the height of the seat induced an earlier hip full extension, which is possibly related to the earlier offset of the muscle potentials during forward rotation of the crank. Practically, increasing the height of the seat is considered for individuals with TFA who suffer from limited hip and knee flexion range of motion, but we should also consider lowering the seat for easier pedaling at the end of the downstroke phase.

The effects of the seat-tube angles can be divided into that of the anteroposterior pelvic tilt and that of the arch-like transition of the seat. The anterior pelvic tilt increased the hip flexion without affecting the knee kinematics, which resulted in increased muscle potential of the proximal adductors and decreased muscle potential of the primary extensors and abductors (Figure 4A). A post-hoc comparison between the results of the seat-tube angle and the pelvic tilt suggested that the arch-like transition of the seat had little effect on the joint kinematics and muscle potential. Therefore, manipulating the pelvic tilt instead of the seat-tube angle could be useful for adjusting the hip flexion angles and the balance of muscle potentials between the primary extensors and the proximal adductors.

**Figure 4.**
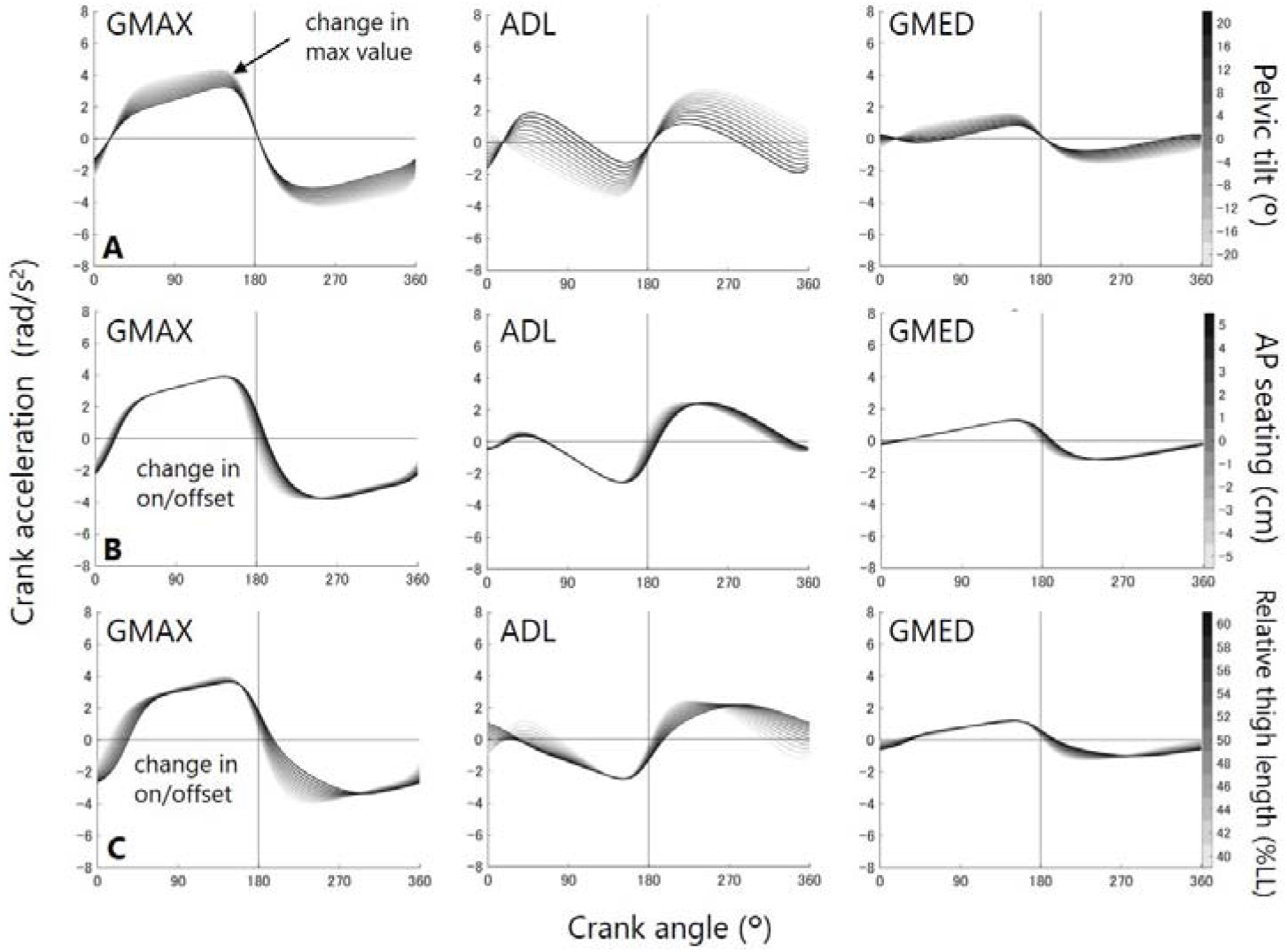
Potential of the residual hip muscles to rotate the crank depending on riding position (A: pelvic tilt, B: anteroposterior [AP] seating position, C: relative thigh length). Abbreviations are the same as Figure 3.

The crank length is a unique variable that can change the range of motion without significantly affecting the muscle potential for pedaling compared with other variables. Choosing a shorter crank for the prosthetic side would be helpful for individuals with TTA^10^, especially for those with limited hip and knee range of motion, even though the pedaling efficiency might be compromised by the reduced length of the moment arm for crank rotation.

The effects of the AP seating position can be divided into that of the seat-tube angle and that of the seat height change. Anterior translation of the seating position increases the seat-tube angle and reduces seat height resulting in combined effects on the pedaling kinematics and the onset/offset of muscle potentials for forward rotation of the crank. The AP seating position influences the knee kinematics, especially during the downstroke phase and the hip kinematics during the upstroke phase. These characteristics could help when adjusting the knee kinematics during the downstroke phase is preferred. In addition to these theoretical effects, the anterior seating position might be an option to help avoid contact between the prosthetic socket and the saddle, a frequent concern for cycling after TFA^4^.

The relative thigh length is not usually adjusted to the bicycle for each individual with TFA, but this information about its effect on pedaling could be useful as the relative thigh length depends on the residual limb length, which varies between individuals. Increasing the thigh length reduces the hip peak flexion and the range of motion without significantly changing the knee kinematics, which is possibly related to the reduction in muscle potential during the downstroke phase for proximal adductors.

This simulation study has several limitations. First, this study focused mainly on the lower limb kinematics and the muscle potential (i.e., effects of unit muscle force) for pedaling. This means that the force-length and the force-velocity relationship of the muscles, and the individual muscle forces required for pedaling were not reflected in the results. For example, the crank length and AP seating position had relatively smaller effects on the muscle potentials, but they altered the pedaling kinematics, which could influence the pedaling performance. Second, the volume and line of action of the residual muscles could differ according to the residual limb length^23^ or amputation surgery^35^. We should consider the altered geometry and force capacity of biarticular muscles (i.e., hamstrings in this study) when interpreting the current simulation results. Finally, while simulation results are suggestive, experimental evaluation of subjects with TFA is necessary to confirm the current simulation findings. However, we believe the pedaling kinematics generated in this study would not be significantly different from the real one because pedaling with a passive transfemoral prosthesis is a constrained movement with a fewer number of active joints compared with pedaling with an intact limb. Possible contributors we did not consider for the current model were pelvic motion and out-of-plane prosthetic limb motion, which would not be as large as hip and knee flexion. The reaction force from the pedal can retract the residual limb from the socket during pedaling, which should be kept in mind especially when using a suspension system without rigid fixation. This can be evaluated by pedaling force measurement in future experimental studies.

In conclusion, the seat height, crank length, and pelvic tilt are the primary candidates for bicycle fitting considering their adjustability and their simple effects on the joint kinematics and muscle potential. The seat-tube angle and AP seating position practically have the combined effects of the other variables and thus can be reserved for fine-tuning after gross fitting of the bicycle. Considering the effects of relative thigh length could help as it also affects hip kinematics and muscle potentials, although it is usually not considered for adjustment.

## Acknowledgment

This study was supported by JSPS KAKENHI grant number JP18K17744.

