## Supplementary material for "Effects of bicycle geometry and riding position on the potential of residual limb muscles to pedaling with a transfemoral prosthesis: a computer simulation study": suppl

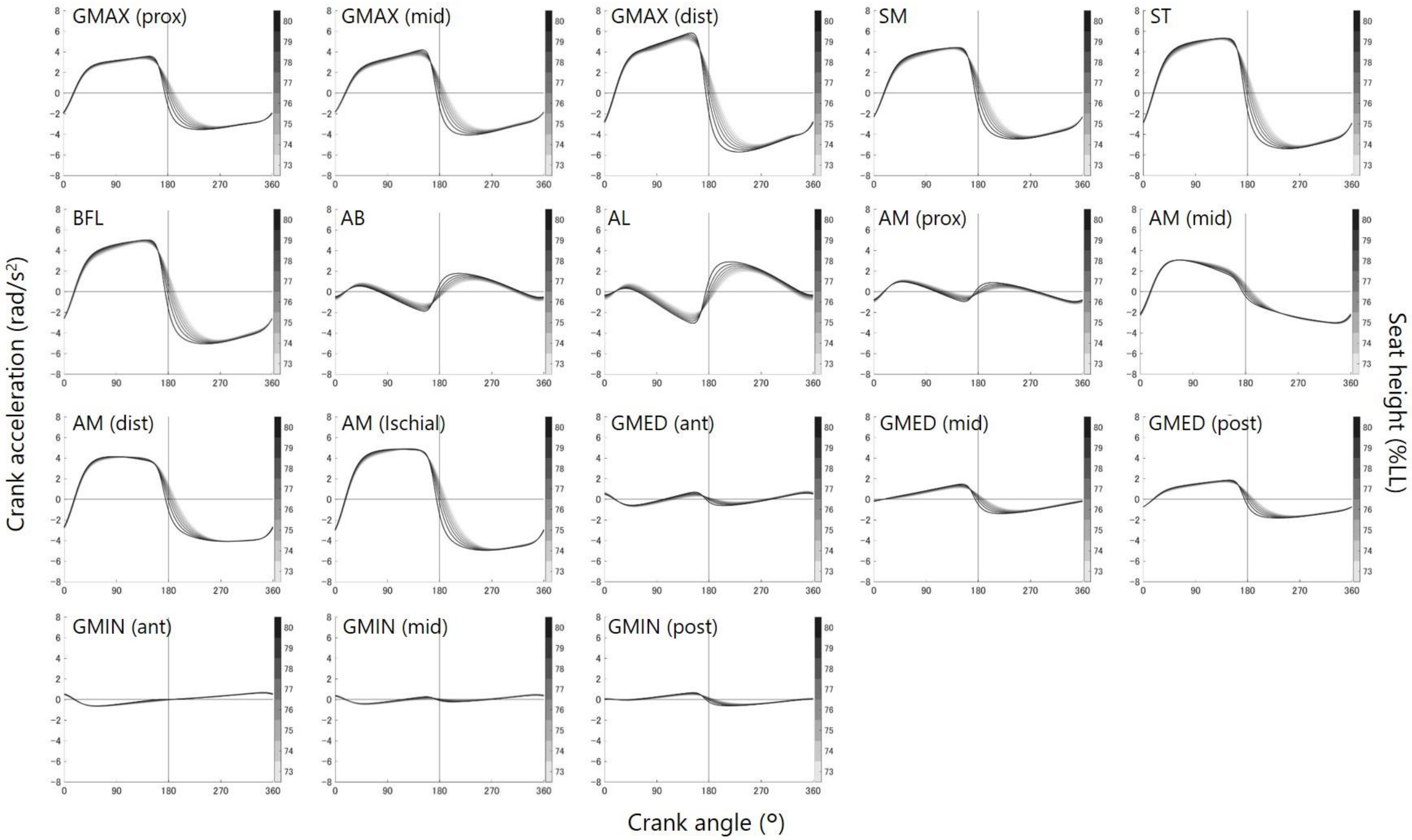


Supplementary figure 1. Individual muscle potential to rotate the crank. The color density of the line indicates the seat height. Abbreviations: GMAX: gluteus maximus, SM: semimembranosus, ST: semitendinosus, BFL: the long head of biceps femoris, AB: adductor brevis, AL: adductor longus, AM: adductor magnus, GMED: gluteus medius, GMIN: gluteus minimus. Some muscles were separated to represent multiple parts: anterior (ant), posterior (post), proximal (prox), middle (mid), distal (dist), and ischiocondylar (isch).

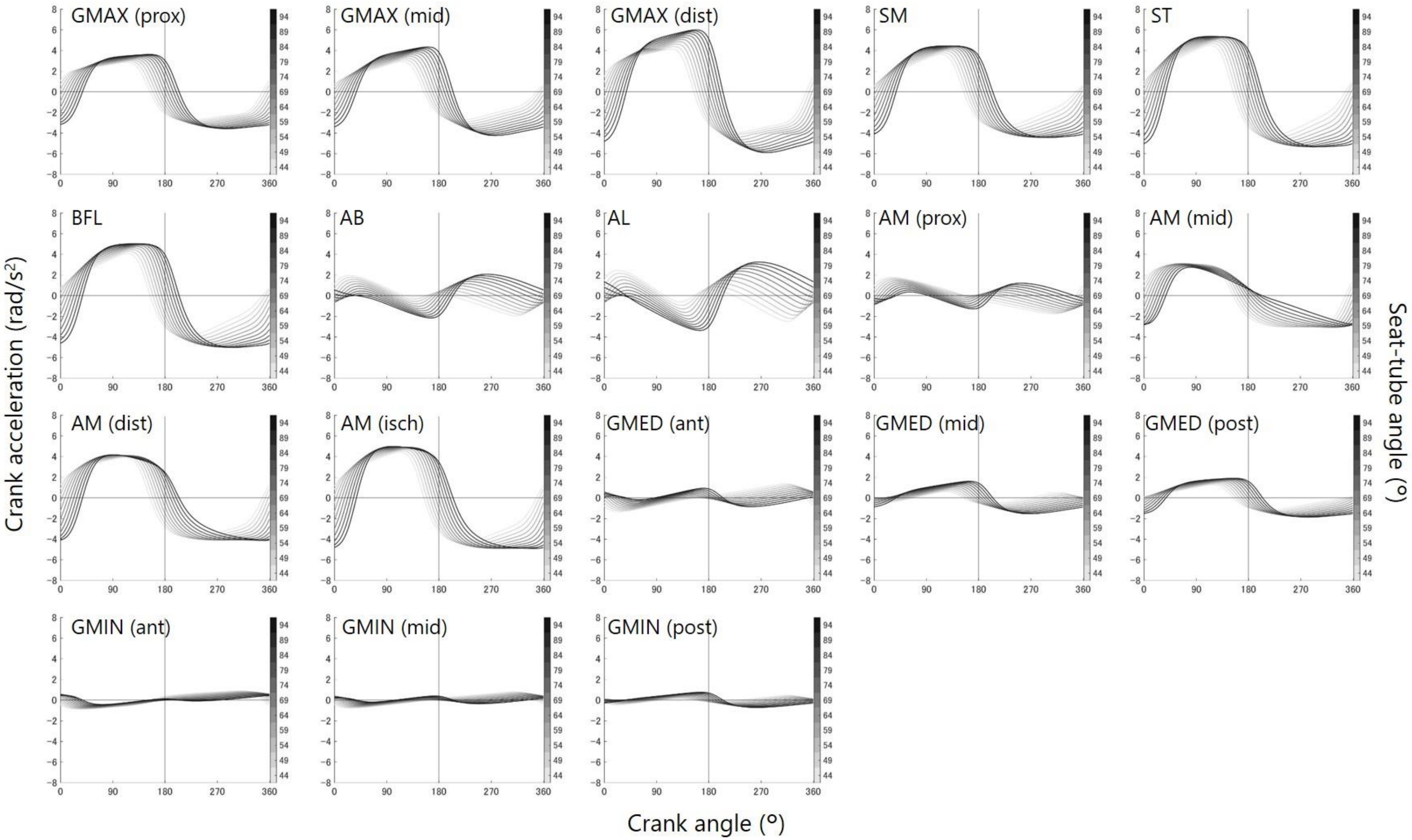


Supplementary figure 2. Individual muscle potential to rotate the crank. The color density of the line indicates the seat-tube angle. See Supplementary figure 1 for abbreviations.

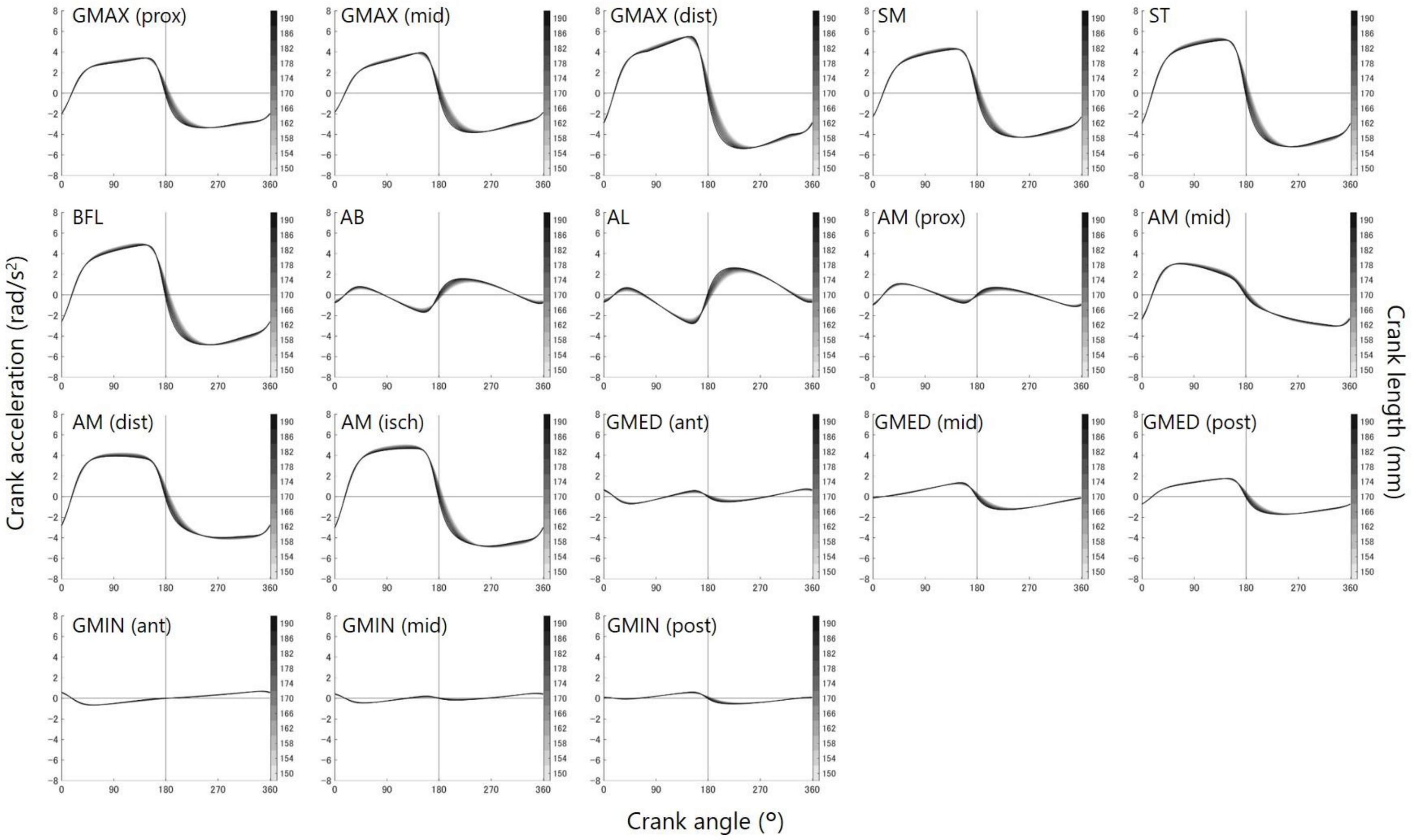


Supplementary figure 3. Individual muscle potential to rotate the crank. The color density of the line indicates the crank length. See Supplementary figure 1 for abbreviations.

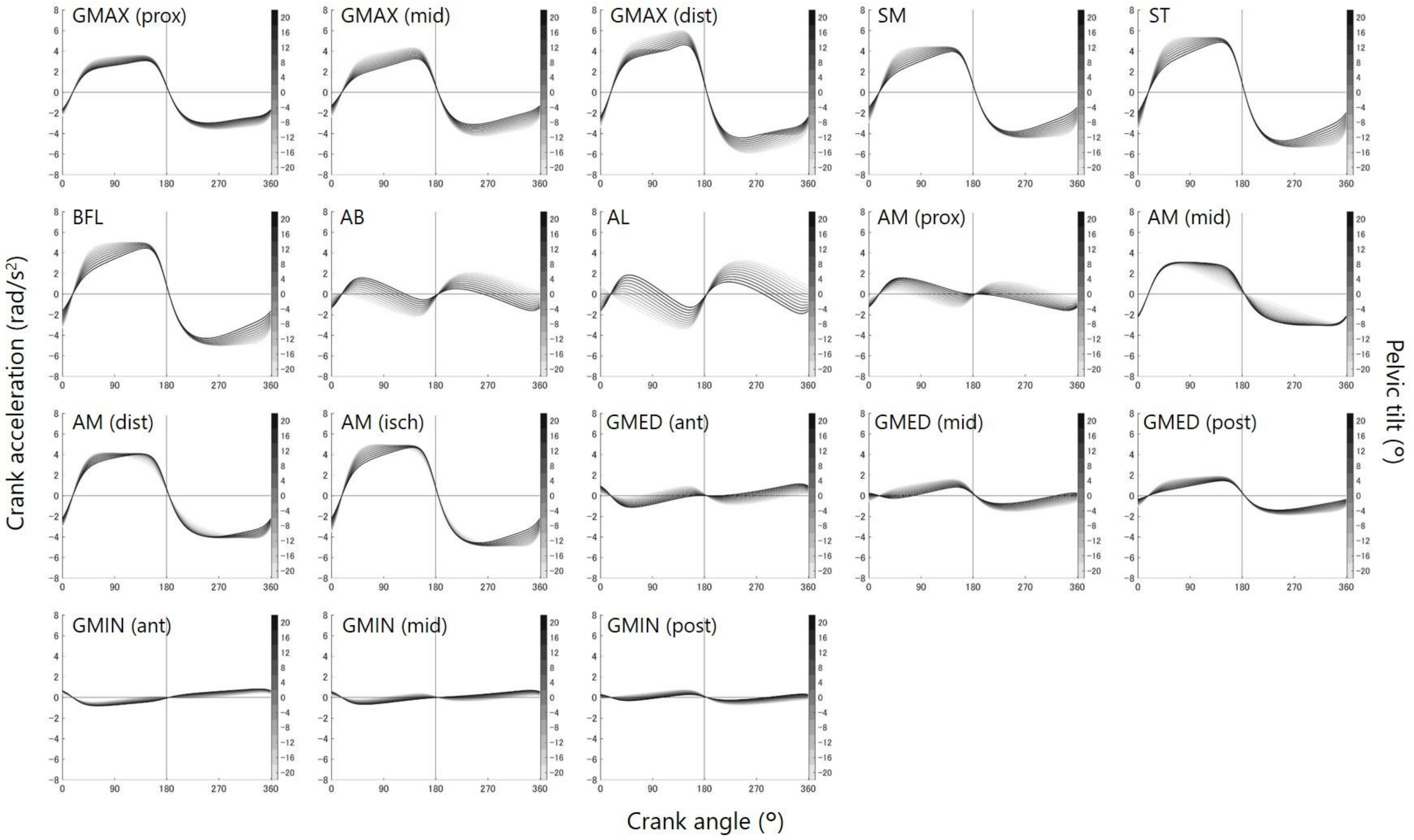


Supplementary figure 4. Individual muscle potential to rotate the crank. The color density of the line indicates the pelvic tilt. See Supplementary figure 1 for abbreviations.

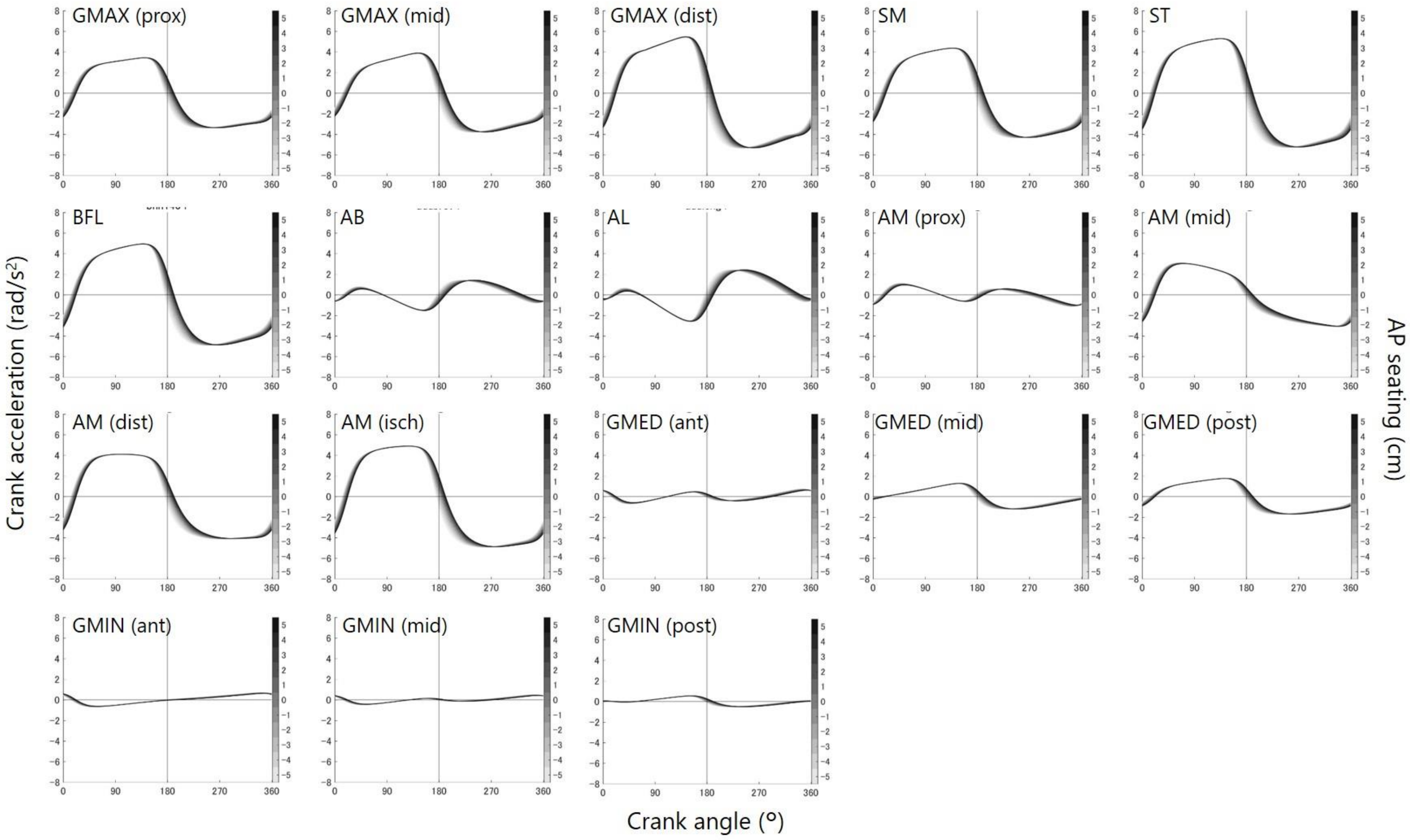


Supplementary figure 5. Individual muscle potential to rotate the crank. The color density of the line indicates the anteroposterior (AP) seating position. See Supplementary figure 1 for abbreviations.

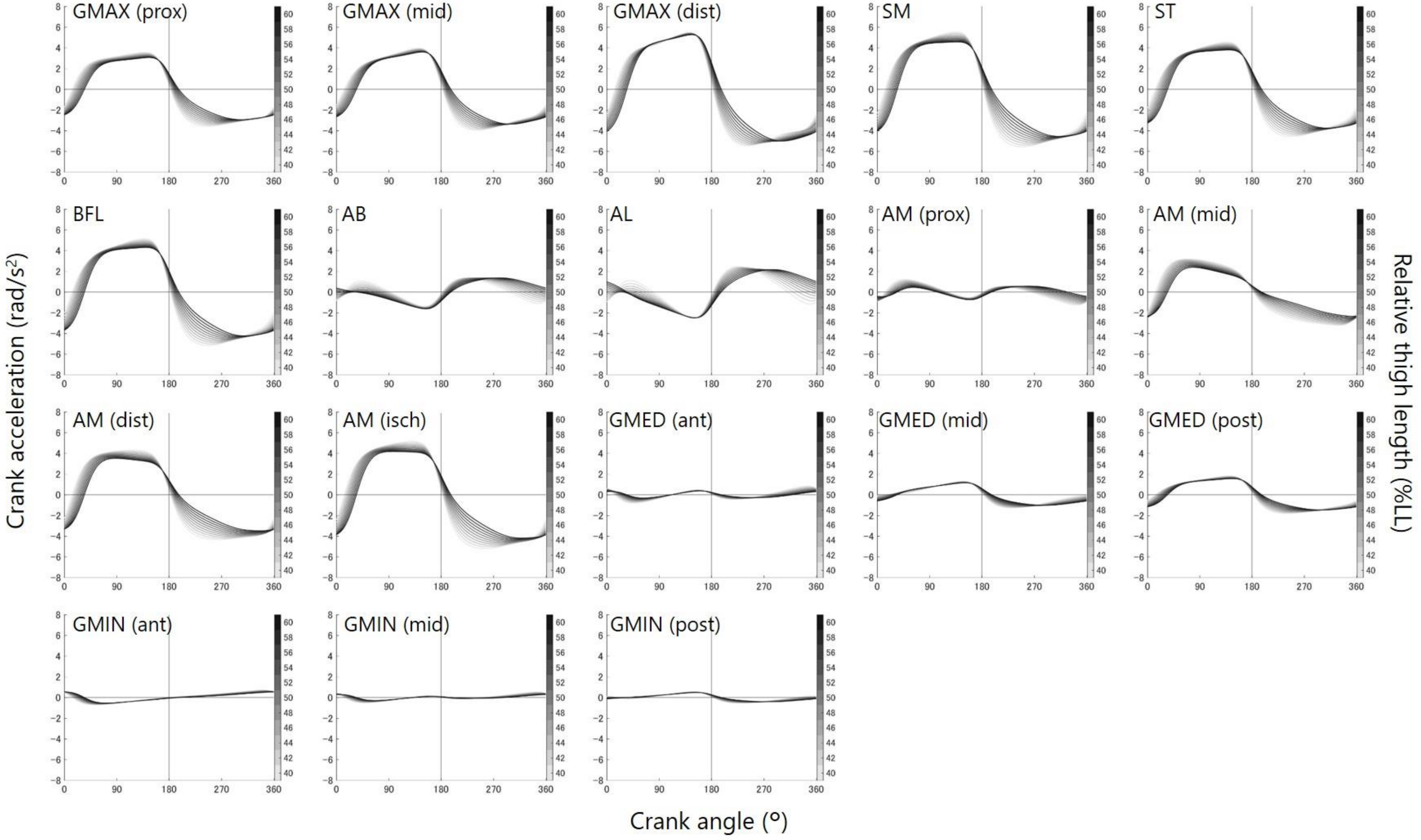


Supplementary figure 6. Individual muscle potential to rotate the crank. The color density of the line indicates the thigh length relative to the entire leg (% leg length [%LL]). See Supplementary figure 1 for abbreviations.
